# Identification of host factors for Rift Valley Fever Phlebovirus

**DOI:** 10.1101/2023.09.28.559935

**Authors:** Velmurugan Balaraman, Sabarish V. Indran, Yonghai Li, David A. Meekins, Laxmi U.M.R. Jakkula, Heidi Liu, Micheal P. Hays, Jayme A. Souza-Neto, Natasha N. Gaudreault, Philip R. Hardwidge, William C. Wilson, Friedemann Weber, Juergen A. Richt

## Abstract

**Background:** Rift Valley fever phlebovirus (RVFV) is a zoonotic pathogen that causes Rift Valley fever (RVF) in livestock and humans. Currently, there is no licensed human vaccine or antiviral drug to control RVF. Although multiple species of animals and humans are vulnerable to RVFV infection, host factors affecting susceptibility are not well understood.

**Methodology:** To identify the host factors or genes essential for RVFV replication, we conducted a CRISPR-Cas9 knock-out screen in human A549 cells. We then validated the putative genes using siRNA-mediated knockdowns and CRISPR-Cas9-mediated knockout studies, respectively. The role of a candidate gene in the virus replication cycle was assessed by measuring intracellular viral RNA accumulation, and the virus titers by plaque assay or TCID_50_ assay.

**Findings:** We identified approximately 900 genes with potential involvement in RVFV infection and replication. Further evaluation of the effect of six genes on viral replication using siRNA-mediated knockdowns found that silencing two genes (WDR7 and LRP1) significantly impaired RVFV replication. For further analysis, we focused on the *WDR7* gene since the role of *LRP1* in RVFV replication was previously described in detail. Knock-out A549 cell lines were generated and used to dissect the effect of *WRD7* on RVFV and another bunyavirus, La Crosse encephalitis virus (LACV). We observed significant effects of *WDR7* knock-out cells on both intracellular RVFV RNA levels and viral titers. At the intracellular RNA level, *WRD7* affected RVFV replication at a later phase of its replication cycle (24h) when compared to LACV which was affected an earlier replication phase (12h).

**Conclusion:** In summary, we have identified *WDR7* as an essential host factor for the replication of two relevant bunyaviruses, RVFV and LACV. Future studies will investigate the mechanistic role by which *WDR7* facilitates Phlebovirus replication.

**Authors Summary:** Rift Valley fever phlebovirus is a high consequence pathogen that infects multiple animal species and also humans. Currently, there are no control measures available to treat RVF in humans and to prevent the incursion of Rift Valley fever virus into non-endemic countries. RVFV poses a significant threat to animal and human health in countries where it is endemic. RVFV replication depends on the host’s machinery to complete its replication cycle. Therefore, one way to control virus replication is to disrupt the interaction between the virus and the host proteins important for replication. In this study, we identified a host factor, the WDR7 gene, that is critical for RVFV replication. The identification of this host factor is important as it can potentially lead to the development of antiviral strategies to control Rift Valley fever in both humans and animals.

## Introduction

Rift Valley fever phlebovirus (RVFV) is a mosquito-borne, segmented RNA virus that belongs to the family *Phenuiviridae,* genus *Phlebovirus.* RVFV was first isolated and characterized in the Rift Valley of Kenya in 1931[1] and is the causative agent of Rift Valley fever (RVF). It is endemic throughout sub-Saharan Africa [2], the Arabian peninsula (Saudi Arabia, Yemen) and Mayotte [3,4]. RVFV can be naturally transmitted to and cause disease in several species of animals such as cattle, sheep, goats and camels [5–7]. We have recently shown that white-tailed deer are highly susceptible to experimental infection with RVFV [8]. RVF in livestock is characterized by abortion storms in pregnant ewes and pregnant cattle and causes 100% mortality in newborn animals [5–7]. In humans, RVFV infection may be subclinical or cause mild flu-like symptoms and sometimes severe disease with hepatitis, retinitis and encephalitis [9,10] with a small number of cases being lethal [11]. RVFV can infect and replicate in a multitude of cell-lines (e.g., neurons, epithelial cells, etc.) from different animal species such as frogs, pigs, elk, mule deer, pronghorn, reptiles, among others [12–17]; this highlights the potential for the virus to infect a wide variety of animal species.

RVFV is mainly transmitted by infected mosquitoes (*Culex* and *Aedes*), by direct contact with infected animal secretions and exudates [18,19], or by aerosol exposure [20]. Currently, there are no FDA approved therapeutic drugs or licensed vaccines available to control RVF in humans [20]. There is a real risk of introduction of arboviruses such as RVFV to non-endemic countries, such as Europe, Asia, and North America [21], where competent vector mosquito species (e.g., *Culex* and *Aedes*) are present [18,22–24]. Therefore, RVFV poses a global threat to the health of livestock and humans, and to animal trade and commerce [24].

The successful development of antiviral therapies requires the detailed knowledge of viral protein function or of host factors that support virus replication [25]. RVFV enters cells by receptor-mediated endocytosis and releases its nucleocapsid after fusion of virus-endosomal membranes. After completion of replication, the viral particles assemble and bud from the Golgi apparatus [26]. Like many other RNA viruses, RVFV depends on various host factors to complete its replication cycle [27–29]. Several groups have conducted exploratory studies aimed to find host factors or co-factors that might play a role in RVFV replication [27,29–37]. Notably, other researchers have shown that *LRP1* [29,37], heparin sulfate [33] play essential roles in cell entry of RVFV. Furthermore, exogenous administration of the LRP1 inhibitor mRAP_D3_ protected mice from infection with a virulent strain of RVFV [29]. Devignot et al. 2023 reported that a *LRP1* gene knock-out in Huh cells significantly affected intracellular RVFV RNA accumulation [37]. Bracci et al. 2022 found that *UBR4* depletion affects RVFV production and virus titer in mammalian and mosquito cells [36]. Although these studies have identified host factors in mouse cells associated with RVFV replication, none of the host factors were able to completely abolish productive RVFV infection in gene-edited knock-out cells. This indicates that RVFV interacts with different host factors to complete its replication cycle exploiting multiple redundant cellular pathways. Our studies had the following aims: 1) to identify unique host factors that could significantly affect RVFV infection and replication, 2) to identify host factors that could be used as a potential drug target; and 3) to identify host factors that are conserved between different host species. To this end, a genome-wide CRISPR-Cas9 knock-out (GeCKO) screen in human A549 cells infected with the RVFV MP-12 vaccine strain was performed in order to identify host factors essential for RVFV infection and replication. We identified the *WDR7* gene as a critical host factor that plays a role in the late phase of RVFV replication. In addition, the *WDR7* gene also plays a role in the replication of another bunyavirus, the La Crosse encephalitis virus.

## Methods

### Cells

A549 cells (ATCC^®^ CCL-185™, American Type Culture Collection, Mansassas, VA, USA) were cultured in F-12 medium (ATCC, Mansassas, VA, USA), supplemented with 10% fetal bovine serum (FBS, R&D Systems, Minneapolis, MN, USA) and 1% penicillin-streptomycin solution (ThermoFischer Scientific, Waltham, MA, USA). The Vero-MARU cell line is a clone of Vero cells obtained from the Middle America Research Unit. The Vero-MARU, MRC-5 (ATCC® CCL-171™), and Vero E6 (ATCC® CRL-1586™) cell lines were cultured in Dulbecco’s Modified Eagle’s Medium (DMEM, Corning, New York, N.Y, USA), supplemented with 5% FBS (R&D Systems, USA) and 1% penicillin-streptomycin solution (ThermoFischer Scientific, USA). All mammalian cells were maintained at 37°C under a 5% CO_2_ atmosphere. The *Aedes albopictus* larva (C6/36, ATCC® CRL-1660™) cells were maintained at 28^°^C, and cultured in L-15 medium (ATCC, USA), supplemented with 10% insect cell culture tested FBS (IFBS, catalog. no: F4135, Sigma-Aldrich, St. Louis, MO, USA), 10% tryptose phosphate broth (TPB, catalog. no: T9157, Sigma-Aldrich, USA), and 1% penicillin-streptomycin solution (ThermoFischer Scientific, USA).

### Virus Strains

The RVFV MP-12 vaccine strain provided by US Army Medical Research Institute for Infectious Diseases [38] was propagated in MRC-5 cells; the RVFV Kenya 128B-15 virulent strain was provided by R. Bowen, Colorado State University with authorization from B. Miller, Centers for Disease Control, Fort Collins, CO [39] was grown in C6/36 cells. La Crosse Encephalitis virus (LACV), NR-540, was obtained from BEI resources, NIAID, and propagated in Vero E6 cells. RVFV MP-12 and Kenya 128B-15 strains were titered by plaque assay and the LACV by TCID_50_-CPE assay. All the assays involving the pathogenic RVFV Kenya 128B-15 strain was carried out in a BSL3+ facility at Biosecurity Research Institute of Kansas State University.

### Generation of GeCKO-A549 Cell Line and RVFV Screen

The lentiCRISPRv2 library, which targets 19,000 human genes, was obtained from Addgene (catalog number: 1000000048, Addgene, USA). The library contains non-target control sgRNAs, sgRNAs targeting miRNAs, and six unique sgRNAs designed to target each individual human gene. To generate GeCKO-A549 cells, a pooled lentivirus library was created using the lentiCRISPRv2 plasmids, following previously described methods [40, 41]. A puromycin (catalog. no: A1113803, Sigma-Aldrich, USA) cytotoxicity curve was performed on A549 cells, and the puromycin concentration used was determined to be 2 μg/ml medium. Then, transduction efficiency of the lentivirus library on A549 cells was determined as previously described (40,41). Two independently pooled GeCKO-A549 cell lines were generated and subjected to forward genetic screening. Briefly, 80 million GeCKO-A549 cells were subjected to up to three rounds of cytolytic infection with RVFV MP-12 (1 MOI), and the surviving cells were expanded between each round of infection. The gDNAs were extracted from the round 0 (mock-infected), round 1 and 3 virus infections of GeCKO-A549 cells using the midi gDNA extraction kit (Qiagen, Germantown, MD, USA). The sgRNA’s DNA copies were PCR amplified from the extracted gDNAs for next generation sequencing (**Fig 1**). Next generation sequencing was performed using NextSeq (Illumina, USA), and the obtained data were analyzed using MAGeCK software. The ranking of genes were determined using robust ranking aggregation [42].

**Fig 1:**
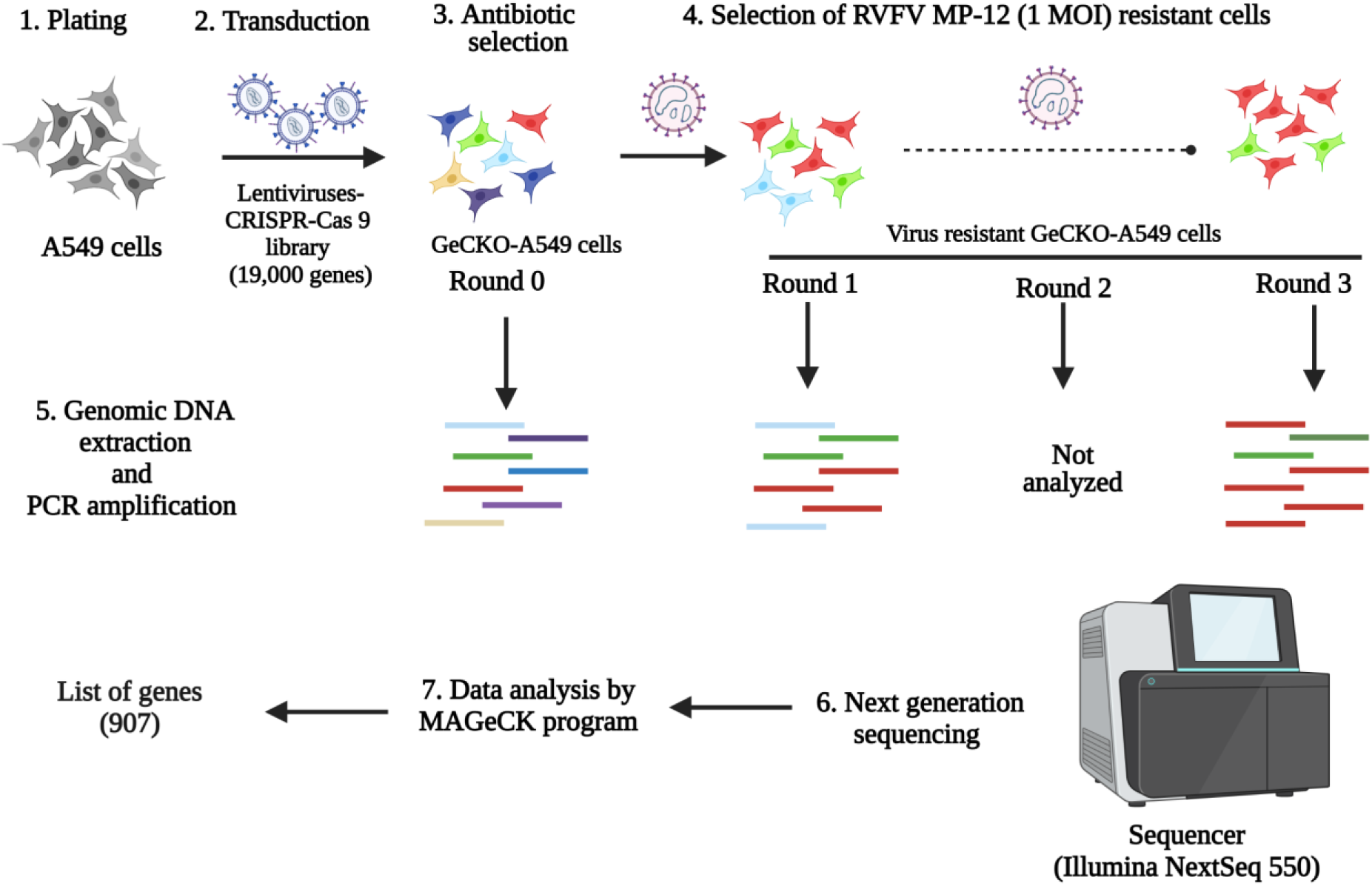
Schematics of GeCKO-A549 cells generation, selection, NGS, and data analysis. A549 cells were transduced with lentivirus-CRISPR-Cas9 library to generate GeCKO-A549 cells. Then, the GeCKO cells were subjected to three rounds of infection with RVFV MP-12 (1 MOI) virus. The genomic DNA of round 0 GeCKO-A549 cells, the round 1, and the round 3 virus resistant GeCKO cells, were sequenced using Illumina NextSeq 550 platform. The output NGS data was analyzed by the MaGeCK program to generate the list of genes involved in RVFV replication.

### siRNA Transfection

Six genes were selected after NGS analysis of the RVFV resistant GeCKO-A549 cells for siRNA gene knock-down studies (**S1 Table**). The gene targets for the siRNAs were as follows: siRNAs: NTC-non-target control catalog. no: D-001206-14-05; WDR7 catalog. no: M-012867-01-0005; LRP1 catalog. no: M-004721-01-0005; EXOC4 catalog. no: M-013068-01-0005; SLC35B2 catalog. no: M-007543-01-0005; and EMC3 catalog. no: M-010715-00-0005); they were commercially purchased (Dharmacon, USA). The positive control siRNA-si46N [43] targeting the RVFV nucleoprotein was obtained from Integrated DNA Technologies (USA). A549 cells were plated in 96-well plates and incubated overnight. The cells were transfected with siRNAs (50nM) using lipofectamine RNAimax reagent (ThermoFischer Scientific, USA). Forty-eight hours post-transfection, cells were infected with RVFV MP-12 at 0.1 MOI and the infected cell supernatant was collected at 24 hours post-infection. The virus titer of the supernatants was determined by plaque assay on Vero-MARU cells.

### RT-qPCR for Host Gene Expression

To confirm gene knock-down, two step RT-qPCR assays were performed. Briefly, A549 cells were transfected with gene specific siRNAs at 50nM and 48 hours later, the total cellular RNA was extracted. The RNA extraction was performed using the RNAqueous Micro total RNA isolation kit (ThermoFischer Scientific, USA) following the manufacturer’s protocol. Prior to cDNA synthesis, residual gDNA was removed from the extracted RNA using DNAse I enzyme (ThermoFischer Scientific, USA). Then, 400 ng of RNA was used for cDNAs synthesis using the Superscript IV First-Strand Synthesis kit with oligo dT primers (ThermoFischer Scientific, USA) following the manufacturer’s protocol. All RT-qPCR reactions were performed in a CFX96 Real-Time thermocycler (BioRad, Hercules, CA, USA). The standard real-time qPCR assays were performed using Perfecta Fastmix II (Quanta BioSciences, Beverly, MA, USA) with gene specific primers (**S2 Table**); the glyceraldehyde 3-phosphate dehydrogenase (GADPH) gene was used as an internal control [44]. The percentage gene knock-down was calculated using the 2^−ΔΔC^T method [45].

### Generation of WDR7 Knock-out (KO) cells

Two WDR7 knock-out (KO) cell lines and a control non-KO cell line were generated as previously described [41]. WDR7-targeting sgRNAs (sgRNA 1: 5’ GTGACATCCTGTTACGATCG 3’ and sgRNA 5: 5‘AAGATGGCAAGATCGATGCT’3) were applied to generate 2 WDR7 KO cell lines, WDR7 KO cell lines 1 (WDR7 KO 1) and 2 (WDR7 KO 2). The non-KO control cell line (CT) was generated by transduction of the lentiCRISPRv2 vector with the Cas9 backbone without sgRNAs. LentiCRISPRv2 plasmids 1 and 5 containing sgRNAs specific for WDR7 gene were purchased from Genescript, USA. The control and WDR7 sgRNA plasmids were packaged into lentivirus, and the A549 cells were transduced with 0.5 MOI of lentivirus. The transduced cells were kept under puromycin selection and passed three times prior to testing. The gDNA of the two WRD7 KO cell lines were extracted using the DNAeasy kit (Qiagen, Germantown, MD, USA), and the gDNA PCR amplified for NGS analysis. The sequencing was performed using a MiSeq (Illumina, USA). The indel percentage of the KO cell lines were calculated using the python script [41].

### Western Blot Analysis

A549 cells, CT cells, and WDR7 KO 1 and 2 cells at passage 3 were used for western blot analyses. The cell lysates were prepared as previously described [46]. Cell lysates containing 55.0 µg total protein were loaded onto 4–12% Bis-Tris polyacrylamide gels (ThermoFischer Scientific, USA), and transferred onto a polyvinylidene difluoride (PVDF) membrane using a Trans-Blot Turbo Transfer Pack (BioRad, USA).

The membrane was blocked using 5% skim milk, and then incubated with a primary polyclonal antibody against WDR7 (diluted 1:500, catalog. no: sab2109026, Sigma-Aldrich, USA) or β-actin (diluted 1:5000, catalog. no: ab20272, Abcam, USA) for 1 h at room temperature. The membrane was then incubated with horseradish peroxidase (HRP)-conjugated polyclonal goat anti-rabbit immunoglobulin (diluted 1:1000, catalog. no: 31460, ThermoFischer Scientific, USA). The target proteins were detected using Super Signal West Femto Maximum Sensitivity Substrate according to the manufacturer’s protocol (catalog. no: 34095, ThermoFischer Scientific, USA). The images were taken using a ChemiDoc MP Imaging System (BioRad,USA).

### Testing of WDR7 KO cells for Virus Replication

The non-knock-out control (CT) and WDR7 KO cell lines were seeded onto 96-well plates and allowed to incubate overnight. Afterwards, the cells were infected with either RVFV MP-12, RVFV Kenya 128B-15, or La Crosse encephalitis virus (LACV) at 0.1 MOI, and the cell supernatants were collected at 6-, 12-, 24-, or 48-hours post-infection (h pi). The titer of collected supernatants was determined using plaque assay (RVFV) or TCID_50_-CPE (LACV) assays.

### Intracellular Viral RNA Accumulation Assay

The viral RNA accumulation was determined at various time points (0, 2, 5 and 24 hours) post-infection (h pi) as previously described [37,47]. The CT and WDR7 KO 1 cells were plated in 6-well plates. Twenty-four hours later, cells were infected with RVFV MP-12 or LACV at a MOI of 0.1 for one hour (h) at 0^°^C to allow virus attachment and entry. For the 0 h infection, immediately after infection, the cells were washed thrice with 1x phosphate buffered saline (PBS [pH=7.2-7.6], catalog. no: P4417, Sigma-Aldrich, St. Louis, MO, USA), lysed in 350 RLT buffer (Qiagen, Germantown, MD, USA), and then stored at -80^°^C till further use. For the post infection time points, the cells were washed once with 1x PBS after the initial 1 hour of incubation, and then incubated with 2 mL of pre-warmed fresh medium. At 2 h pi, cells were first trypsinized and collected into microcentrifuge tubes. Then, the trypsinized cells were washed three times with 1x PBS by centrifugation at 10,000 g for 5 min. The cell pellets were lysed in RLT buffer and stored at -80^°^C till further use. For the 5- and 24-hour time points, the cells were washed once with 1x PBS, and lysed in RLT buffer for 10 min prior to storage at -80^°^C till further use. The total cellular RNA was extracted using RNeasy Mini kit (Qiagen, Germantown, MD, USA). One-step RT-qPCR assays were performed using q-script XLT (2x) Master mix (Quanta BioSciences, Beverly, MA, USA) with virus gene specific primers and probes (**S3 Table**); the phosphoglycerate kinase (PGK1) gene was used as an internal housekeeping control gene [44]. Respective gene expressions were calculated using 2^−ΔΔC^T method [48].

### Plaque Assay

Vero-MARU cells were seeded in 12- or 24-well plates and incubated at 37°C and 5% CO_2_ overnight. After overnight incubation, cells were infected with RVFV for one hour and then the medium was replaced with overlay of 1% methylcellulose-2x MEM (ThermoFischer Scientific, USA),10% FBS, 2% antibiotics/antimycotic. The cells were incubated for 5-7 days and then stained and fixed with 5% crystal violet fixative solution. The plaques were counted, and the titer was expressed as pfu/ml.

### TCID_50_-CPE Assay

Vero E6 cells were seeded in 96-well plates one day prior to infection. Ten-fold serial dilutions of LACV were prepared in 96-well plates in DMEM supplemented with 5%FBS and 1% antibiotics/antimycotic. The diluted viral suspensions were then added onto Vero E6 cells. Three to four days post infection, the cells were visually observed under microscope for CPE and the titer was calculated using the Spearman-Karber method [49].

### Statistical Analysis

Statistical analysis performed in this study is described in the figure legends. All the statistical tests were carried out using GraphPad Prism version 9.3.0.

## Results

### Identification of host factors involved in RVFV replication

To identify genes potentially involved in RVFV replication, we performed CRISPR-Cas9 knock-out screens in A549 cells. The A549 type II alveolar human cell line was selected for the screen because it is susceptible to RVFV and can be easily transduced with the human GeCKO library. The GeCKO-A549 cells were subjected to three rounds of infection with the RVFV MP-12 vaccine strain to select for resistance to RVFV infection to identify key host factors that are required for virus replication. Extensive cytopathic effect (CPE) was observed during the first round of infection. Surviving cells were re-infected and the CPE was much less extensive during the second and third round of infection. To assess the susceptibility of the round 3 GeCKO-A549 cells after three rounds of RVFV infection, virus growth kinetic assays were performed. A significant difference in MP-12 virus titers were observed between the round 0 and the round 3 GeCKO-A549 cells at 24-and 48-hours post-infection (hpi) (**S1 Fig**), indicating that the round 3 GeCKO-A549 cells had acquired resistance to RVFV infection. Next, genes involved in RVFV replication were determined by analyzing the NGS data from round 0, round 1 and round 3 GeCKO-A549 cells. Our analysis of the round 3 GeCKO-A549 cells revealed that 907 genes (p-value <0.05) seem to be involved in RVFV MP-12 replication (**S1 Data**). For further analysis, we selected the six top genes significantly enriched in round 3 GeCKO-A549 cells: LRP1, SLC35B2, EMC3, WDR7, EXOC4 and CT47A1 (**S1 Data**). We did not investigate the other top two genes, ART3 and CEBPD (**S1 Data**), as they were associated with essential cellular functions.

### Validation of genes from the pooled GeCKO-A549 cell screen

To assess the effect of the six top genes enriched in the round 3 GeCKO-A549 cells on RVFV replication, we used siRNA-mediated gene silencing (gene knock down) in A549 cells. Gene knock-down was confirmed by respective RT-qPCR assays and the average reduction of gene expression ranged from approximately 55% to 90% (**S2 Fig**). After gene knock-down, the cells were infected with RVFV MP-12 virus at 0.1 MOI for 24 hrs, supernatants were harvested, and extracellular virus titer determined by plaque assay. There was an average of 56% or 42% reduction in virus titer upon *WDR7* and *LRP1* knock-down, respectively, compared to non-target control (NTC) siRNA targeting the firefly luciferase mRNA (**Fig 2**). The positive control siRNA, siRNA-si46N, targets the N protein gene of RVFV, and caused a reduction of approximately 96 % in virus titer compared to the negative control group. We observed no significant effect on virus titers following the knock-down of the other 4 selected top genes, *EXOC4, CT47AL1, EMC3* and *SLC35B2* (**Fig 2**). These results demonstrate that the knock-down of *WDR7* and *LRP1* significantly impaired RVFV replication. Given that the role of *LPR1* gene in RVFV replication has been recently demonstrated [29,37], we focused our further analysis on the newly discovered putative RVFV host factor *WDR7*.

**Fig 2:**
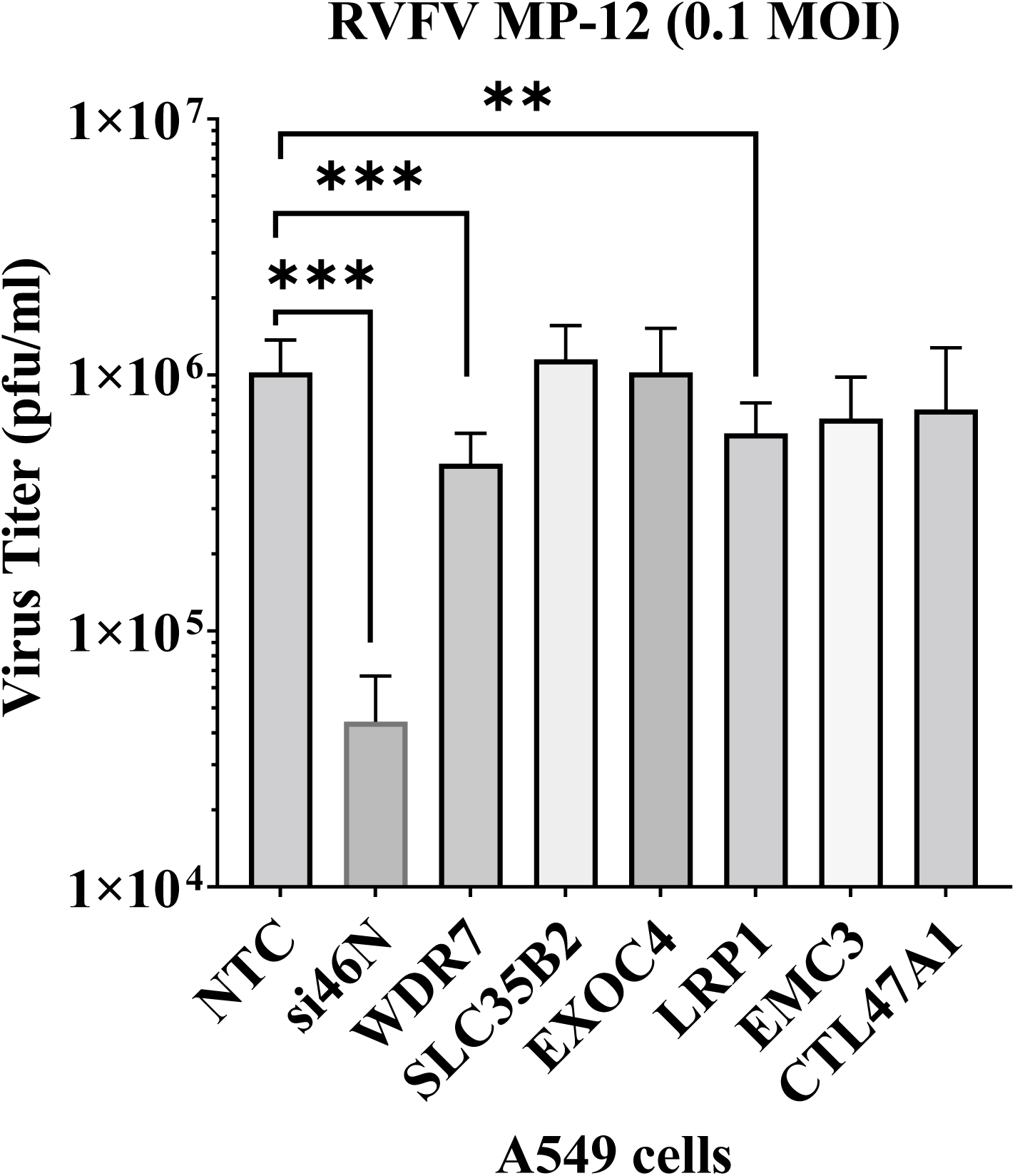
Validation of gene hits by siRNA gene knock-down study. A549 cells were transfected with 50 nM of siRNAs. At 48 hours post-transfection, the cells were infected with RVFV MP-12 virus at 0.1 MOI. At 24 hours post-infection, the supernatant was collected and titered by plaque assay. NTC-non-target control siRNA, si46N-anti-RVFV siRNA, WDR7-, SLC35B2-, EXOC4-, LRP1-, EMC3-, CTL47A1-gene specific siRNAs. Each bar represents the average virus titer (pfu/ml) along with the corresponding standard deviation. Statistical analysis was done on two independent experiments with four replicates for each, using Mann-Whitney U independent Student’s t-test (** p-value < 0.005, *** p-value <0.001).

### Generation and characterization of knock-out cells

To investigate the role of *WRD7* in the RVFV replication cycle, we employed highly enriched sgRNAs targeting the *WDR7* gene to generate two knock-out A 549 cell lines: WDR7 KO line #1 and WDR7 KO line #2. The established WRD7 knock-out cells were analyzed by NGS sequencing, which confirmed *indels* in nearly 100% of the WDR7 KO cells (99% and 98%, respectively, for the two WDR7 KO cell lines #1 and #2, (S1 Table)). There was also a significant decrease in WDR7 protein expression in the WDR7 KO cell lines as compared to the control and non-transduced A549 cells (**Fig. 3A**). However, we noted the presence of faint WDR7 band in both WDR7 KO cells.

**Fig 3:**
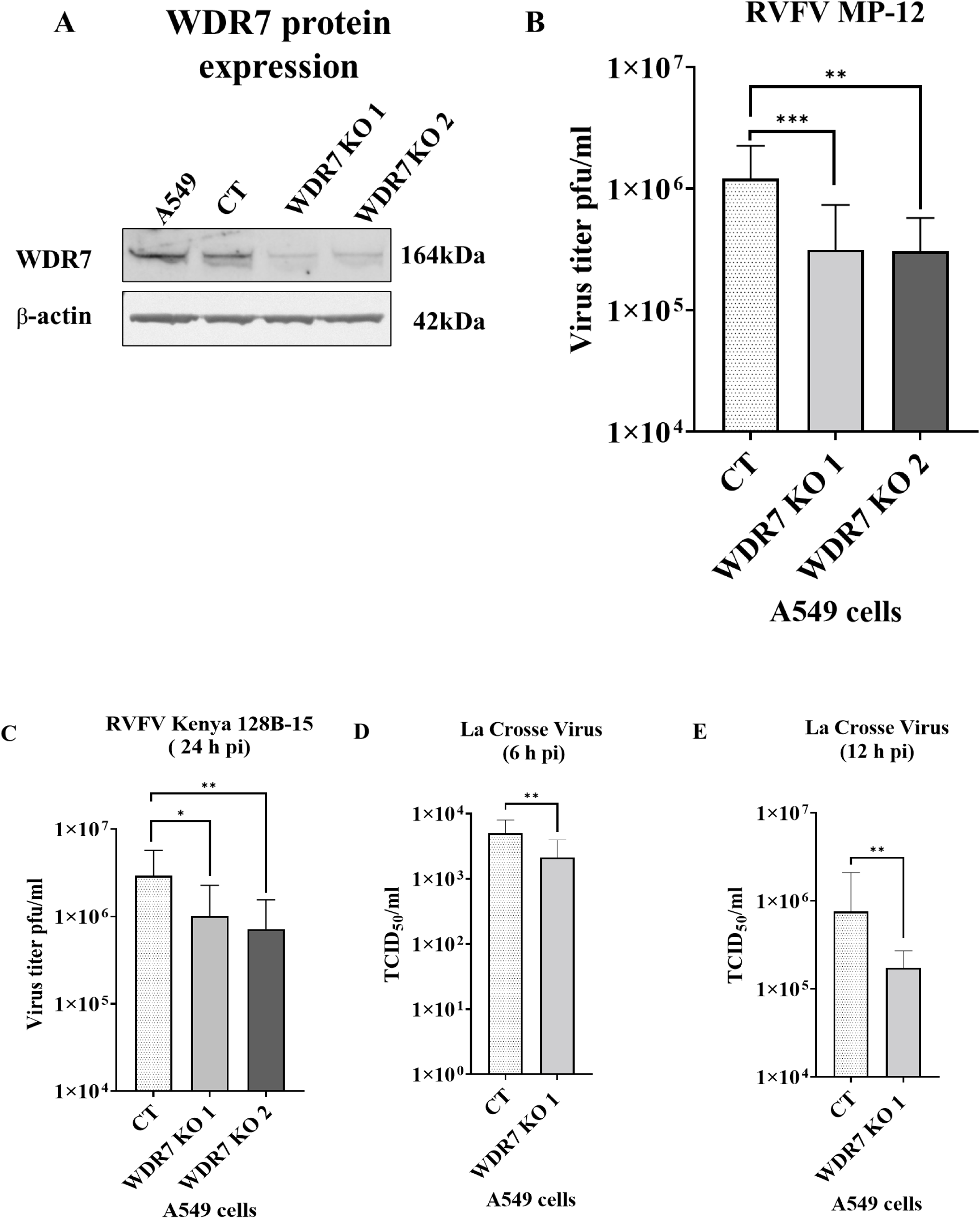
Effect of WDR7 gene knock-out (KO) on virus production of bunyaviruses. (**A**) A549 cells, CT (non-knock-out control) cells, and WDR7 gene KO cell lines #1 and #2 were analyzed for WDR7 protein expression by western blot using a WDR7-specific polyclonal antibody. (**B, C, D & E**) CT cells and WDR7 KO A549 cells were infected with RVFV MP-12 vaccine strain, (**B**) with the wild-type RVFV Kenya 128B-15 strain, (**C**) or with La Crosse encephalitis virus (**D, E**) at 0.1 MOI. Supernatant was collected at 6, 12 or 24 h post infection (h pi) and titered by plaque assay (RVFV) or TCID_50_-CPE assay (LACV). RVFV MP-12 testing on CT A549 cells, WDR7 KO lines #1 or #2, and NTC-non-target control cells involved three to five independent experiments with three to four technical replicates each. RVFV Kenya 128B-15 testing involved independent experiments with three technical replicates each. LACV testing was performed in two independent experiments with eight technical replicates each. Statistical analysis was done using Mann-Whitney U independent Student’s t-test (* p-value < 0.05, ** p-value < 0.005, *** p-value <0.001).

To ensure the authenticity of the A 549 control cells, we sequenced the *WDR7* gene at the target site and found the *WDR7* gene is not mutated in the CT cells (**S1 sequence file**); also, the WDR7 protein expression in CT cells was at a similar level as in the non-transduced A549 cells (**Fig 3A**). Moreover, the CT cells showed comparable levels of virus replication to the non-transduced wild-type A549 cells (**S3E Fig**). Additionally, cell viability did not differ significantly between the *WDR7* KO cell lines #1 and #2, and the CT cell lines, neither prior to or after RVFV MP-12 infection (**S3A-S3D Fig**).

### Effect of WDR7 gene knock-out on RVFV and LACV infection

Next, we infected the two WDR7 KO cell lines with the RVFV MP-12 strain at 0.1 MOI and determined the extracellular virus titers by plaque assay. The *WRD7* gene KO resulted in a significant reduction of approximately 74% in virus titer compared to CT cells at 24h post infection (**Fig 3B**), while no difference in virus titers were observed at 48h post infection (**S4A Fig**). We then evaluated the effect of the *WDR7* gene KO on the virulent RVFV strain Kenya 128B-15. Our results showed an average reduction of RVFV Kenya 128B-15 titers of 66% and 75% in WDR7 KO cell lines 1 and 2, respectively, compared to the CT cells (**Fig 3C**). Taken together, these findings support the results obtained using the siRNA knock-down assays and confirm a critical role of the *WDR7* gene on the RVFV replication cycle.

In addition, we evaluated if the *WDR7* gene plays a role in the infection cycle of other bunyaviruses. For this purpose, we used La Crosse encephalitis virus (LACV) and infected the CT and WDR7 KOA549 cell #1 line with LACV; the cell supernatant was collected at various time points post-infection and the virus titer determined by TCID_50_-CPE assay. The results showed an average reduction in LACV titer of 57 % and 77% at 6 h pi and 12 h pi, respectively, in the WDR7 KO #1 cell line compared to the control CT cells **(Fig 3D and 3E**). However, at 24 h pi, the reduction in virus titer was approximately 39% but did not reach statistical significance (**S4B Fig**). Overall, these findings highlight the importance of the *WDR7* gene also in LACV replication.

### WDR7 gene KO impairs RVFV and LACV intracellular RNA accumulation

To investigate the role of *WDR7* in the RVFV and LACV replication cycle, we quantified intracellular viral RNA accumulation at 0 hour(s) post-infection (h pi; attachment phase), 2 h pi (entry phase), 5 h pi (replication phase) and 24 h pi (late phase of replication) using previously established protocols [37,47]. At 0, 2 and 5 h pi, there was no significant difference in RVFV RNA accumulation between the control and WDR7 KO cells (**Fig 4A**). However, at 24 h pi, we observed a significant reduction in virus RNA accumulation between the WDR7 KO and control cells (**Fig 4A**). When we infected the WDR7 KO and control cell lines with LACV, we found that WDR7 KO cells had higher levels of LACV RNA accumulation at 0 h pi, i.e. the attachment phase, compared to the control cells (**Fig 4B**). However, at early time points (2 and 5 h pi) and up to 24 h pi, we observed a significant reduction in LACV RNA accumulation in WDR7 KO 1 cells when compared to the control cells (**Fig 4B**). These results suggest that *WDR7* disruption affects intracellular viral RNA accumulation primarily at the late phase of the RVFV replication cycle and at an early phase of LACV replication cycle.

**Fig 4:**
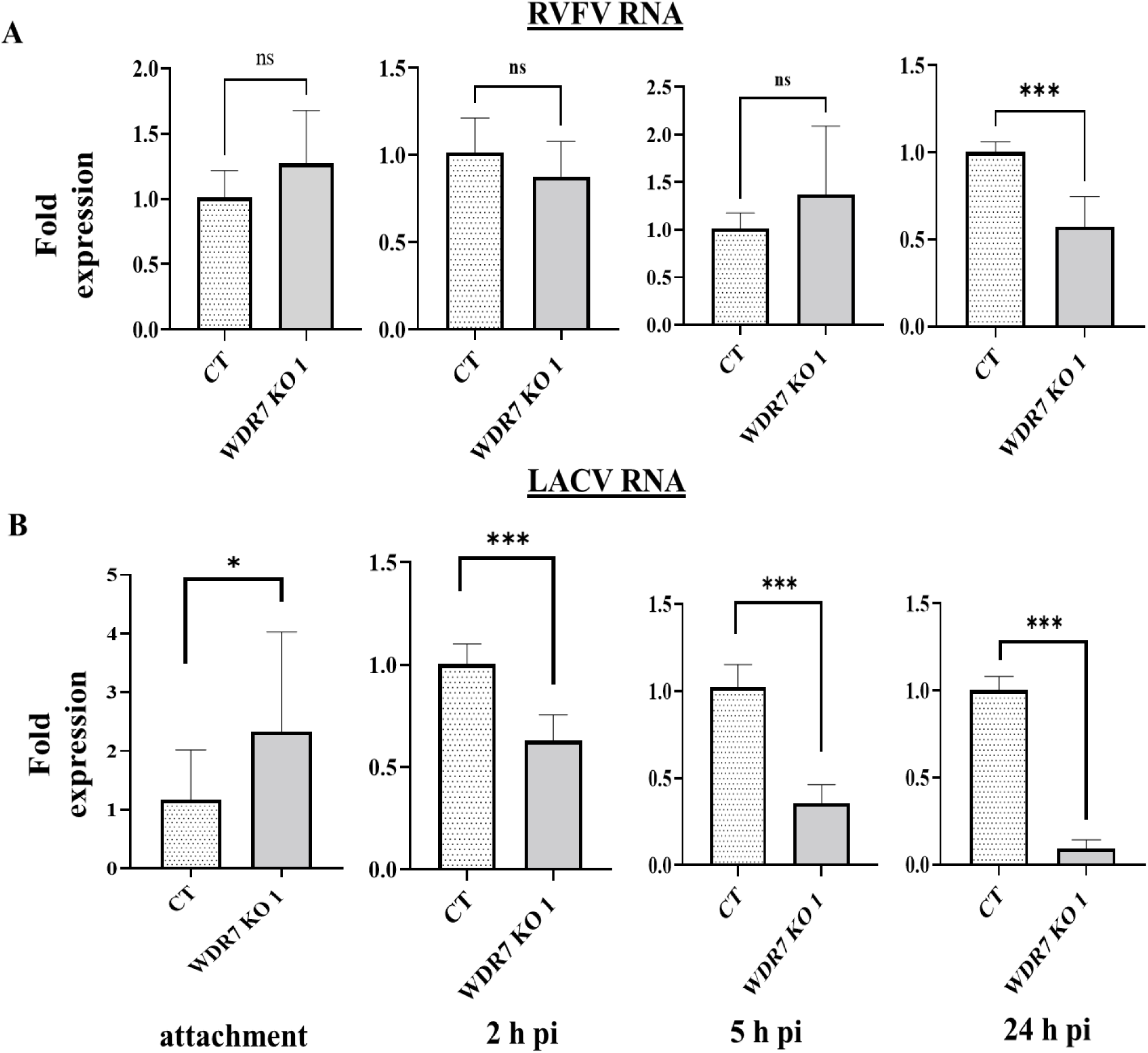
Viral RNA accumulation at various time points post infection in WDR7 knock-out (KO) cells. CT and WDR7 KO #1 cells were infected with the (**A**) RVFV MP-12 vaccine strain or (**B**) the LAC virus at 0.1 MOI. Total cellular RNA was harvested at various hour(s) post-infection (h pi). One-step RT-qPCR was performed to detect the level of viral RNA using the PGK1 gene as an internal control. CT and WDR7 KO #1 cells were utilized. Each bar graph represents the average fold change in viral RNA expression, along with the corresponding standard deviation. Statistical analysis was done on three independent experiments with two to three technical replicates for each, using Mann-Whitney U independent Student’s t-test (* p-value <0.05, *** p-value <0.001, ns, non-significant).

## Discussion

RVFV has a broad cell-tropism and is reported to infect several animal and mosquito species. As discussed previously, RVFV interacts with different host factors in a variety of cell types [29–34,36,37]. We identified *WDR7*, a member of the WD repeat protein family, as a host factor important in the lifecycle of bunyaviruses. We confirmed WDR7 gene knockout through NGS and indel analysis, but western blotting detected a faint band corresponding to WDR7 protein.

This minimal expression could be due to a single guide sgRNA inducing minor double-stranded DNA breaks that resulted in the production of some non-functional protein. We demonstrated that disruption of the WDR7 gene impairs viral RNA accumulation and infectious virus production of two bunyaviruses, RVFV and LACV. However, the exact role of *WDR7* in the replication cycle of these viruses needs further investigation. Previous studies have shown that *WDR7* also plays a significant role in the replication cycle of other RNA viruses such as Dengue, Zika, West Nile virus [50] and influenza A virus [51].

*WDR7* has been associated with V-ATPase, which mediates intracellular vesicle acidification in mouse kidney cells [52], suggesting that *WDR7* could be playing a role in endocytosis or secretory pathways within the virus replication cycle. Here, we demonstrated that *WDR7* affects the late phase of RVFV replication cycle, as shown by the reduction of intracellular viral RNA in *WDR7* KO cells compared to non-KO CT cells at 24 h pi. Combined with the lower levels of infectious RVFV in *WDR7* KO cell supernatants, this might suggest that *WDR7* impacts virus egress and release. In contrast, for LACV, *WDR7* seems to affect virus entry since a significant reduction in both intracellular viral RNA and infectious virus production was found at an early time point post-infection, along with higher levels of virus attachment in KO cells compared to CT cells. This could be due to the fact that the *WDR7* gene KO might affect the expression or function of other host factors involved in virus attachment to the cell surface, or that the knockout of *WDR7* affects the conformation or expression of cell surface molecules needed for attachment.

Interestingly, the effect of the KO of *WDR7* in A549 cells on virus replication appears to diminish at later replication time points for both RVFV and LACV. This pattern is consistent with the findings reported by Bracci et al, (2022), who observed a similar trend in RVFV replication in *UBR4* knock-out cells, with a significant reduction at 24 h pi, but no significant effect at 48 h pi [36]. This suggests that RVFV and LACV have the ability to utilize multiple alternative host factors and pathways to complete its replication cycle. We also observed a significant reduction in LACV viral RNA at later time points, but not in infectious virus production. This result could be attributed to various factors such as a gene knockout effect on late RNA synthesis, increased RNA degradation, or decreased RNA stability in the absence of *WDR7*; all these could affect viral RNA synthesis or RNA stability while virus release or egress was unaffected.

Overall, this study highlights the importance of the *WDR7* gene in bunyavirus replication and suggests that it could be a potential target for the development of antiviral therapies. Further research, including *in vivo* studies using KO mouse models, is needed to fully elucidate the role of *WDR7* in bunyavirus replication.

## Data availability

S1 Data: List of genes enriched. This data will be provided upon request.

S1 Table: Location of the indels in WDR7 gene of Knock-out (KO) cell populations

S2 Table: Primer used for gene expression by qPCR, NGS and sanger sequencing.

S3 Table: Primer and probe list for detection of viral RNA by qPCR.

S1 Fig: RVFV growth kinetics on GeCKO-A549 cells and wild-type A549 cells:

S2 Fig: Confirmation of gene knockdown

S3 Fig: Cell viability of control or gene knockout A549 cells

S4 Fig: Effect of WDR7 gene KO on the replication of RVFV at a later time point

S1 Sequence file: Nucleotide sequence of WDR7 gene-Exon 1-28 in knockout cell line.

## Disclaimer

Mention of trade names or commercial products in this publication is solely for the purpose of providing specific information and does not imply recommendation or endorsement by the U.S. Department of Agriculture. The conclusions in this report are those of the authors and do not necessarily represent the views of the USDA. USDA is an equal opportunity provider and employer.

## Acknowledgements

We are grateful to the staff of KSU Biosecurity Research Institute. We also thank Cassidy Keating for assistance with cell culture and PCR assays. Dane Jasperson, USDA, ARS Center for Grain and Animal Health Research, Insect and Cell Culture Laboratory, has provided C6/36 cells.

## Authors contribution

V.B. Conceptualization, Investigation, Methodology, Validation, Formal Analysis, Writing-Original Draft, Review & Editing; S.V.I. Conceptualization, Investigation, Methodology, Writing-Review & Editing; Y.H.L, D.M, L.U.M.R, H.L, M.H, J.S.N, N.G, P.H, F.W. Investigation, Methodology, Writing-Review & Editing ;W.W, and J.R. Conceptualization, Funding Acquisition, Project Administration, Resources, Supervision, Writing-Review & Editing.

