## Supplemental Figures & Tables for "Identification of host factors for Rift Valley Fever Phlebovirus"

| **Cells** | **Nucleotide sequence-WDR7 gene (5’-3’)** | **Amino acid sequence** |
| --- | --- | --- |
| Predicted nucleotide sequence-WDR7 gene - Exon-2 | **ATG…..ACGATCGTAACAGGATGTCACGAC…** | M…TIVTGCHD… |
| WDR7 KO cell population 1- Exon-2- **‘A’ insertion** | **ATG…..ACGAATCGTAACAGGATGTCACGAC..** | M…TNRNRMSR…  (frame shift mutation-early termination) |
| WDR7 KO cell population 1-Exon-2- **‘T’ deletion** | **ATG…..ACGA_CGTAACAGGATGTCACGAC..** | M…TT*  (Stop Codon) |
| Predicted nucleotide sequence -WDR7 gene - Exon-17 | **ATG………CGAAGATGGCAAGATCGATGCTTGG….** | M…RRWQRCL… |
| WDR7 KO cell population 2- Exon-17- **‘T’ insertion** | **ATG………CGAAGATTGGCAAGATCGATGCTTGG….** | M…RRLARSML…  (frame shift mutation-truncation) |
| WDR7 KO cell population 2- Exon-17- **‘G’ deletion** | **ATG………CGAAGAT_GCAAGATCGATGCTTGG….** | M…RRCKIDAW…  (frame shift mutation-truncation) |

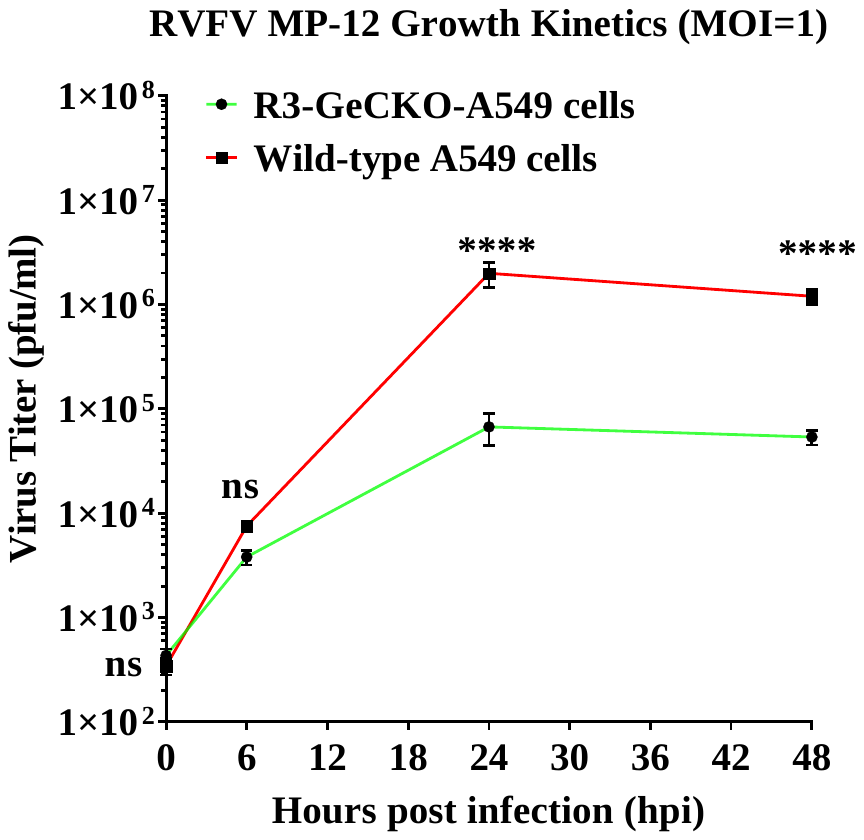

**S1 Fig: RVFV growth kinetics on GeCKO-A549 cells and A549 cells.** The figures illustrate the difference in virus titer between different cell types and time points. Specifically, the virus titer between round 3 MP-12 resistant GeCKO-A549 cells and wild-type A549 cells are compared. Briefly, the cells were infected with RVFV MP-12 virus and the collected supernatant for plaque assay. Statistical analysis was performed on one independent experiment with three replicates, using a 2-way ANOVA sidak’s multiple comparison test (**** p-value <0.0001, ns, non-significant).

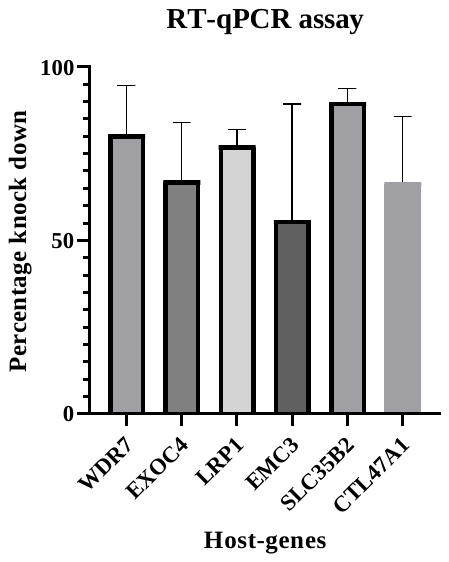

**S2 Fig: Confirmation of gene knockdown.** A549 cells were transfected with 50 nM of siRNAs. Forty-eight hours post-transfection, the total cellular RNA was extracted, and two-step RT-qPCR was performed to determine the percentage of gene knockdown. WDR7- or EXOC4- or LRP1- or EMC3- or SLC35B2- , and CTL47A1- gene specific siRNA transfected A549 cells. Each bar graph represents the average percentage of gene knockdown along with the standard deviation.

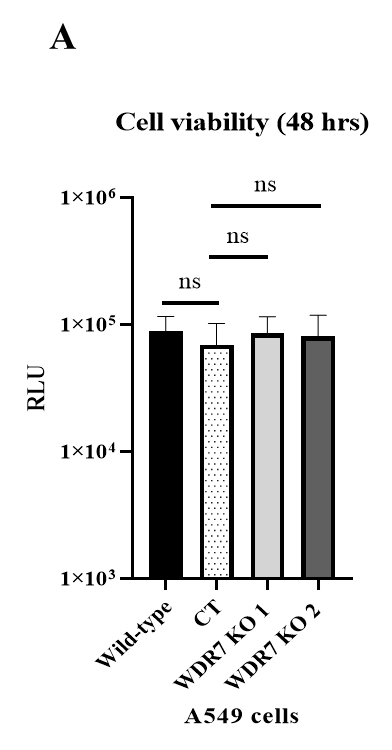

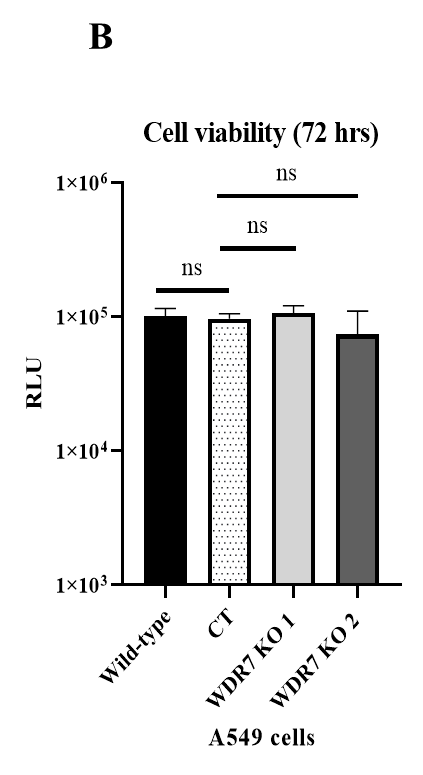

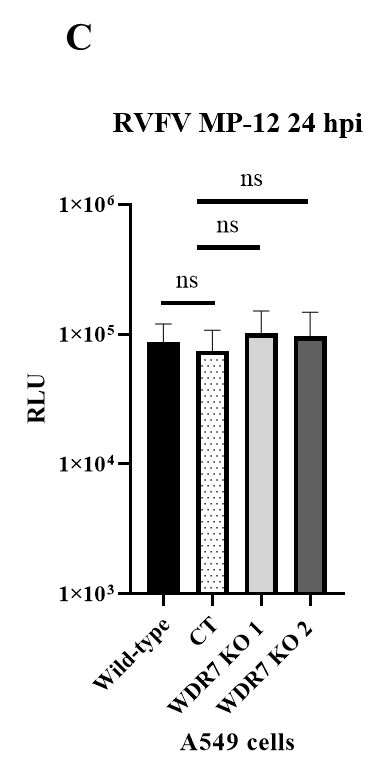

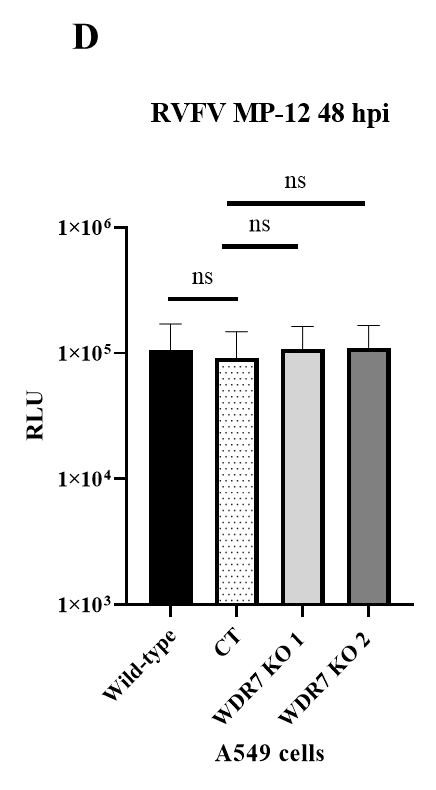

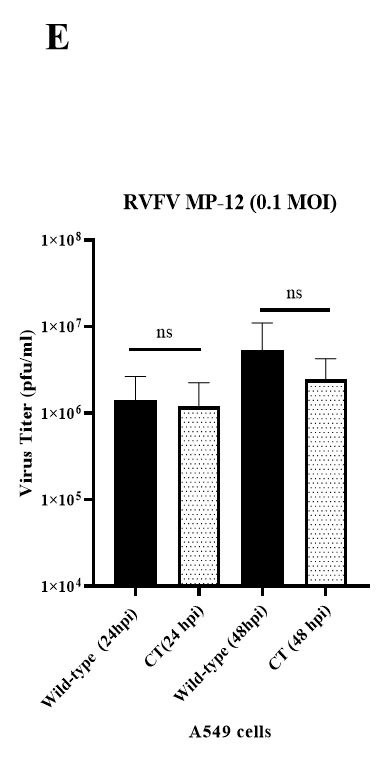

**S3 Fig: Cell viability of control or gene knock outs A549 cells**. (**A & B**) **Uninfected cells**: The A549, CT and WDR7 gene KO cells were plated, and at different time points post-plating, the cell viability was measured by using ADP cell-glow assay. (**C & D**) **Infected cells**: The cell viability of Wild-type, CT and WDR7 gene KO cells were measured after RVFV infection. Each bar graph represents the average Relative Light Unit (RLU; A, B, C D) per sample along with the standard deviation. (**E**) **Comparative infection**: Each bar graph represents the average virus titer (pfu/ml) along with the standard deviation. The level of virus infection between A549 and CT cells was determined by plaque assay. Wild-type A549- non-transduced cells, CT- non-KO A549 cells, WDR7 KO 1 or KO 2- gene knockout cell lines 1 and 2, respectively. Statistical analysis was done on two independent experiments with four technical replicates for each, using Mann-Whitney U independent Student’s t-test (ns, non-significant).

**B**

**A**

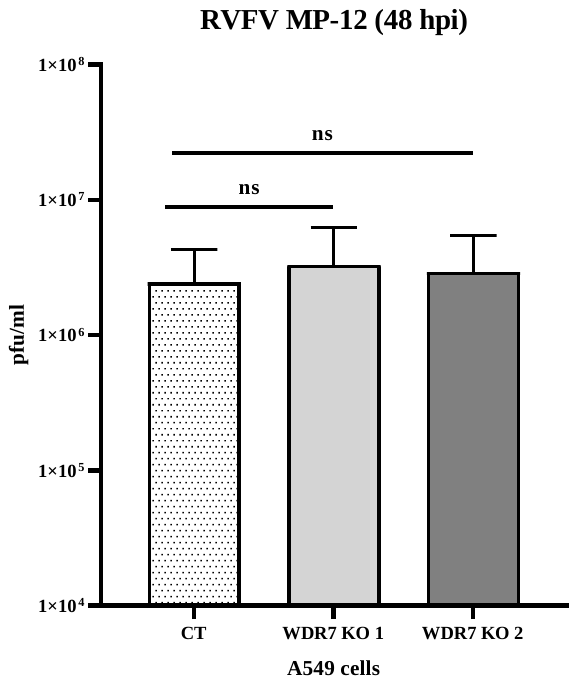

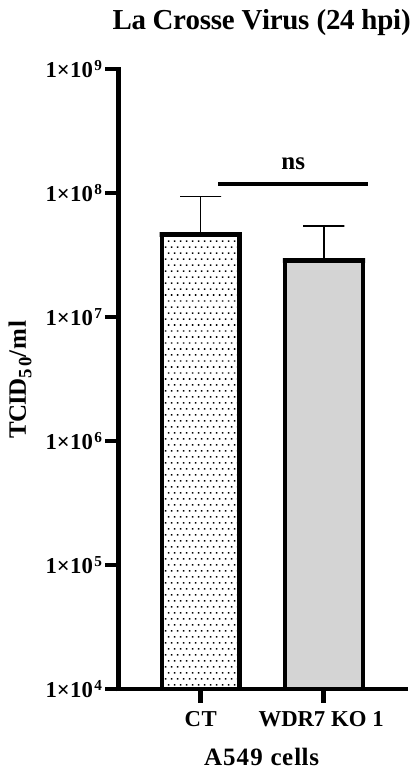

**S4 Fig: Effect of WDR7 gene KO on the replication of RVFV at a later time point.** The CT cells and WDR7 knockout cells were infected with (**A**) RVFV MP-12 or (**B**) LACV virus at 0.1 MOI. At 24- or 48-hours post-infection (hpi), the supernatant was collected and titered by plaque assay (RVFV) or TCID_50_-CPE assay (LACV). CT- non-knockout A549 cells and WDR7 KO 1- WDR7 gene knockout cell line 1. Three independent experiments with three technical replicates each, were performed for RVFV MP-12 testing, and two independent experiments with eight technical replicates each, were performed for LACV. Statistical analysis was done using Mann-Whitney U independent Student’s t-test (ns, non-significant).

**S1 Table: Location of the indels in WDR7 gene of knock-out (KO) cells**. Represent are the predominant indels in the WDR7 gene of WDR7-KO cell lines 1 and 2, respectively. The sgRNA binding sites on WDR7 gene are highlighted in yellow and the indels and the corresponding amino acid changes are highlighted in red. The full nucleotide and amino acid sequences are included in **S1 Sequence file**.

**S2 Table**: **Primers used for gene expression by qPCR, NGS and sanger sequencing**.

| **Gene** | **Primer** | **Sequence** |
| --- | --- | --- |
| WDR7 | WDR7-1 Forward | TTTATGCCACGGACATTACCC |
|  | WDR7-1 Reverse | GGAGCTAATCCAGTCTGGTGATA |
| LRP1 | LRP1-1 Forward | AGCCAGCTATGCACCAACAC |
|  | LRP1-1 Reverse | CCTTGCAGGAGCGGTTATC |
| SLC35B2 | SLC35B2-1 Forward | AGGTGATCCCTGTCATGCTGA |
|  | SLC35B2-1 reverse | CGCTGGATAGCAGAAACATGC |
| EMC3 | EMC3-1 Forward | GTGGTCCTACCCATCGTTATC |
|  | EMC3-1 Reverse | CAGATACTTGTTCCTGGGTGAG |
| EXOC4 | EXOC4-1 Forward | ACAGGTACGTTAATAGTTAATAGCGT |
|  | EXOC4-1 Reverse | TCGTCAATTCTGTGTAGTGCTG |
| GAPDH | GAPDH Forward | TGTAGTTGAGGTCAATGAAGGG |
|  | GAPDH Reverse | ACATCGCTCAGACACCATG |
| CT47A1 | CT47A1-1 Forward | CGTCTGAGACAGACTCTTATTCC |
|  | CT47A1-1 Reverse | TGACCACTGAGGTGGCTA |
| WDR7 Exon 2 | Wsg1-indel-R1-Forward | CTTTCCCTACACGACGCTCTTCCGATCTcCACAAACACAATGGCAGGAAACAG |
|  | Wsg1-indel-R1-Reverse | GACTGGAGTTCAGACGTGTGCTCTTCCGATCT-GGCCCAAAAGCATCATTTGG |
| WDR7 Exon 17 | Wsg5-indel-R1-Forward | CTTTCCCTACACGACGCTCTTCCGATCTtttGACAGGTTGGAGTCAGTTAGCTGC |
|  | Wsg5-indel-R1-Reverse | GACTGGAGTTCAGACGTGTGCTCTTCCGATCT-CAAGTTTACATTTGACCAATGCC |
| NGS- WDR7 Exon 2 | NGS-indel-R2-Forward-1 | AATGATACGGCGACCACCGAGATCTACACTATAGCCTACACTCTTTCCCTACACGACGCTCTTCC |
|  | NGS-indel-R2-Reverse-1 | CAAGCAGAAGACGGCATACGAGATCGAGTAATGTGACTGGAGTTCAGACGTGTGCTCTTC |
| NGS-WDR7 Exon 17 | NGS-indel-R2-Forward-2 | AATGATACGGCGACCACCGAGATCTACACATAGAGGCACACTCTTTCCCTACACGACGCTCTTCC |
|  | NGS-indel-R2-Reverse-2 | CAAGCAGAAGACGGCATACGAGATTCTCCGGAGTGACTGGAGTTCAGACGTGTGCTCTTC |
| Sanger - WDR7 Exon 2 | sWsg1 Forward | TCCCAGCAGGATCTACGCAC |
|  | sWsg1 Reverse | GTCCTCTTGTGTTCGGTGGG |
| Sanger-WDR7 Exon 17 | sWsg5 Forward | GCCACCTAGACCAAGCACC |
|  | sWsg5 Reverse | CGGATAGTGCTGTGTGGAGG |

**S3 Table**: **Primer and probe list for detection of viral RNA by qPCR.**

| **Gene** | **Primer/Probe** | **Sequence** | **Ref.** |
| --- | --- | --- | --- |
| LAC  gene L | qPCR LAC fwd | AGGAAAACTCCTGAGAATATAACTA | (none) |
|  | qPCR LAC rev | GGTATACAAACTGGTGGCGAT |  |
|  | qPCR LAC probe | 6-FAM-CTTAAATTTGAAAATATGTCTAAAATCCAAACATACCCAGGC-BHQ-1 |  |
| RVFV gene L | RVFL-2912fwdGG | TGAAAATTCCTGAGACACATGG | Bird et al., JCM, 2007, 45:11(3506-13) |
|  | RVFL-2981revAC | ACTTCCTTGCATCATCTGATG |  |
|  | RVFL-probe-2950 | 6-FAM-CAATGTAAGGGGCCTGTGTGGACTTGTG-BHQ1 |  |
| PGK1 | PGK1 Forward | GCCACTTGCTGTGCCAAATG | Ori gene |
|  | PGK1 Reverse | CCCAGGAAGGACTTTACCTT |  |
